# Mitochondrial copper and phosphate transporter specificity was defined early in the evolution of eukaryotes

**DOI:** 10.1101/2020.11.02.365692

**Authors:** Xinyu Zhu, Aren Boulet, Katherine M. Buckley, Casey B. Phillips, Micah G. Gammon, Laura E. Oldfather, Stanley A. Moore, Scot C. Leary, Paul A. Cobine

## Abstract

Mitochondrial carrier family (MCF/SLC25) proteins are selective transporters that maintain the mitochondrial metabolome. Here we combine computational, biochemical and phenotypic approaches to understand substrate selectivity of SLC25A3. In mammals, SLC25A3 transports both copper and phosphate, yet in *Saccharomyces cerevisiae* the transport of these substrates is partitioned across two paralogs: PIC2, which transports copper, and MIR1, which transports phosphate. To understand whether the ancestral state of this transporter was a single promiscuous transporter that duplicated and gained selectivity, we explored the evolutionary relationships of PIC2 and MIR1 orthologs across the eukaryotic tree of life. Phylogenetic analyses reveal that PIC2-like and MIR1-like orthologs are present in all major eukaryotic supergroups, indicating that the gene duplication that created these paralogs occurred early in eukaryotic evolution. Frequent lineage-specific gene duplications and losses suggest that substrate specificity may be evolutionarily labile. To link this phylogenetic signal to protein function and resolve the residues involved in substrate selection, we used structural modelling and site-directed mutagenesis to identify PIC2 residues involved in copper and phosphate transport activities. Based on these analyses, we generated a Leu175Ala variant of mouse SLC25A3 that retains the ability to transport copper, but not phosphate, and rescues the cytochrome *c* oxidase defect in *SLC25A3* knockout cells. Taken together, this work uses an evolutionary framework to uncover amino acids involved in substrate recognition by MCF proteins responsible for copper and phosphate transport.

## Introduction

Mitochondrial carrier family (MCF/SLC25) proteins comprise the largest family of mitochondrial inner membrane (IM) proteins and are responsible for transporting numerous substrates including Krebs cycle intermediates, nucleoside di- and triphosphates for energy metabolism and nucleotide replication, amino acids for degradation or maintenance of the urea cycle, and essential metals such as copper (Cu) and iron (1, 2). Structurally, MCF transporters consist of a conserved fold with three repeats that contain two transmembrane helices connected by a short α-helical loop (3, 4). The repeated structural elements and variable copy numbers across eukaryotic phyla (53 in humans and 30 in yeast) suggest that this complex gene family has arisen through multiple duplication events followed by neofunctionalization as substrate needs changed. From an evolutionary perspective, one hypothesis is that protein families with multiple substrates (e.g., enzymes and transporters) arose as generalists that duplicated to evolve specificity over time (5, 6). However, the evolutionary history of the MCF/SLC25 family with respect to substrate specificity remains largely unexplored.

Our current mechanistic understanding of MCF activity is based on *in vitro* transport assays, phenotypic observations made in mutant cells and structures of the ADP-ATP carrier (4, 7). This MCF transporter adopts two conformational states: the cytoplasmic or c-state which is open to the intermembrane space (IMS), and the matrix or m-state which is open to the matrix (8, 9). All MCFs have six transmembrane helices with conserved motifs that allow for formation of salt bridges and the close packing of helices that are critical to the mechanism of transport (4).

Cu is required in mitochondria for the stability and activity of the IM-embedded enzyme cytochrome *c* oxidase (COX) and the IMS-localized superoxide dismutase. The Cu used in the assembly of these enzymes comes from a pool in the mitochondrial matrix (10). We previously identified PIC2 as a mitochondrial Cu transporter in *Saccharomyces cerevisiae* (11). Mutant yeast strains lacking *PIC2* (*pic2Δ*) are deficient in COX activity and have lower mitochondrial Cu levels than isogenic wild-type strains (11). Although PIC2 has also been implicated in phosphate transport (12-15), the major phosphate transporting MCF in yeast is MIR1 (13, 16). *PIC2* expression can complement *mir1Δ* phenotypes and mitochondria from *mir1Δpic2Δ* yeast strains regain phosphate transport activity when PIC2 is overexpressed (12), suggesting that phosphate can be a PIC2 substrate. However, it is unlikely that this transport activity is physiologically relevant under normal conditions as *PIC2* deletion does not result in phosphate deficiency phenotypes in yeast. Based on these findings, we predict that while yeast PIC2 and MIR1 have specialized to transport specific substrates, PIC2 retains some level of promiscuity for Cu and phosphate.

In contrast, humans express a single PIC2/MIR1 paralog, SLC25A3, which serves as the major mitochondrial transporter of both Cu and phosphate (17, 18). Cells lacking *SLC25A3* exhibit a Cu-dependent COX assembly defect (17). Additionally, SLC25A3 transports Cu when recombinantly expressed and reconstituted in liposomes or when heterologously expressed in *Lactococcus lactis* (17). Similarly, both phenotypic and biochemical assays confirm that SLC25A3 is the major phosphate transporter in mammalian mitochondria (14, 18, 19).

These findings highlight a major unanswered question in our understanding of MCFs. Specifically, what differences enable the transport of single versus multiple substrates? Using newly available phylogenomic data from diverse lineages that span the major eukaryotic supergroups, we used an evolutionary framework to infer residues in the PIC2-MIR1 MCF subfamily that likely mediate substrate selection and transport. By coupling phylogenetic analyses with biochemical assays, we have uncovered residues required for transport of Cu and phosphate. Further, we demonstrate that Cu transport to the mitochondrial matrix is directly responsible for the COX deficiency observed in cells lacking *SLC25A3*.

## Results

### MIR1 does not transport Cu

To determine if MIR1 can transport Cu in addition to phosphate, we exploited the fact that MCF proteins insert into the cytoplasmic membrane of *L. lactis* in an active state and that Cu transport activity in this system can be detected by growth arrest in the presence of silver (Ag^+^) (Fig. 1A) (11, 20). This assay was also used to assess phosphate transport by quantifying the growth rates of *L. lactis* strains expressing MCF genes in the presence of the toxic phosphate mimetic arsenate (AsO_4_^3-^). In the presence of 80 µM Ag^+^, the growth of *L. lactis* expressing PIC2, but not MIR1 or an empty vector (EV), was significantly inhibited (Fig 1B). In contrast, the growth of *L. lactis* expressing MIR1 or PIC2 was inhibited to the same extent when cultured in 1.6 mM AsO_4_^3-^relative to a control strain harboring the EV (Fig. 1C). These data show that, in *L. lactis*, MIR1 is capable of transporting the phosphate mimetic AsO_4_^3-^but not the Cu mimetic Ag^+^.

**Figure 1:**
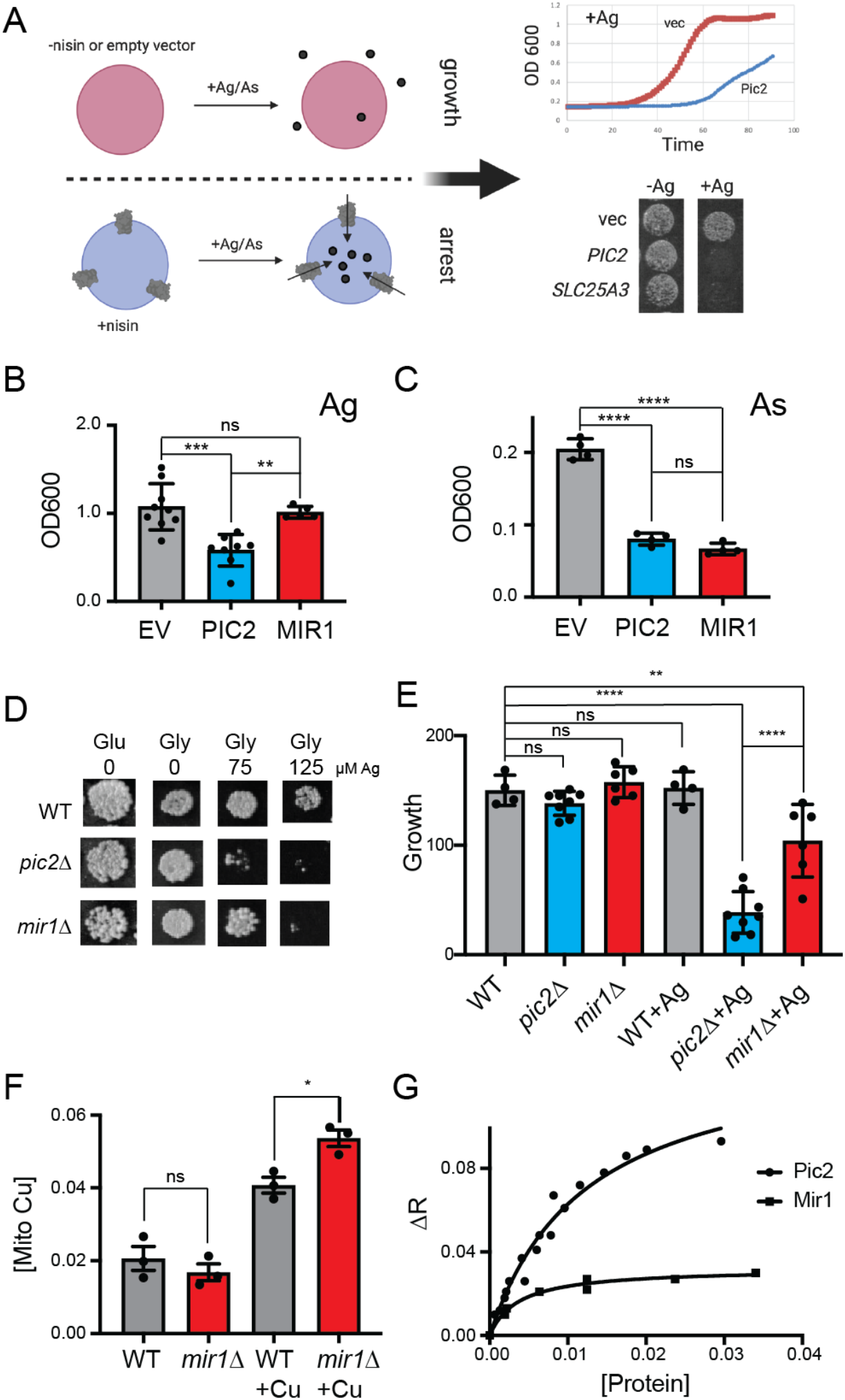
*S. cerevisiae* MIR1 does not transport Cu. *A)* Schematic representation of the *L. lactis* expression system used to quantify transport characteristics. Survival is determined by the growth rate in liquid culture or by visual inspection of cells grown on agar plates containing Ag^+^ or AsO_4_^3-^in the presence of the inducer nisin. *B)* Quantification of the growth of *L. lactis* expressing empty vector (EV), *S. cerevisiae* PIC2 or *S. cerevisiae* MIR1 after 12 hours in 80 µM Ag^+^ containing media (n>5). *C)* Quantification of the growth of *L. lactis* expressing EV, PIC2 or MIR1 after 12 hours in 1.6 mM AsO_4_^3-^containing media (n=5). *D)* Wild-type (WT), *pic2Δ* or *mir1Δ* yeast grown in rich medium with a fermentable (Glu: glucose) or a non-fermentable (Glycerol: Gly) carbon source in the absence (0) or presence of Ag^+^ (75 or 125 µM). All strains were spotted on media as a 10^−3^ dilution of OD^600^ of 1. *E)* Densitometry measurements of serial dilutions (10, 10^2^, 10^3^, 10^4^) of cells in *D)* on Glu, Gly and Gly plus 75µM Ag (WT n=4, *pic2Δ* n=8, *mir1Δ* n=6). *F)* Cu content of purified intact mitochondria from *mir1*Δ cells assayed by ICP-OES and compared with that of parental WT cells. Both strains were grown in YP medium with glucose as a carbon source containing 10 μM BCS or 100 μM Cu (+Cu) (n=3). *G)* Fluorescence anisotropy (FA) of CuL (Ex320, Em400) upon the addition of reconstituted PIC2 or MIR1 in proteoliposomes prepared from extracted egg-yolk lipids. Control FA of equal quantity of lipids without protein added was subtracted from each data point. Protein concentrations were determined by Bradford assay, and curves are fit with a nonlinear regression that assumes a single binding site. In all panels, data are plotted as the mean ± standard deviation and a one-way ANOVA was used for statistical analysis; ns- not statistically significant, * P<0.05, ** P<0.01, ***, P<0.001, ****P<0.0001.

Consistent with our previous results (11), we find that the growth of yeast lacking *PIC2* is severely compromised on a non-fermentable carbon source in the presence of 75 µM Ag^+^ (Fig. 1D,E). In contrast, yeast lacking *MIR1* only exhibited a mild growth defect relative to the isogenic wild-type strain at this Ag^+^ concentration (Fig 1D,E). Exposure to 125 µM Ag^+^ led to a growth defect in both *mir1Δ* and *pic2Δ* yeast but not in the isogenic, wild-type (WT) strain (Fig 1D). To further establish that MIR1 is incapable of Cu transport activity, we quantified mitochondrial Cu levels by inductively coupled optical emission spectroscopy. Cu levels in mitochondria from *mir1Δ* yeast cells were similar to those isolated from WT cells (Fig. 1F). In yeast mitochondria, Cu is stably bound by a fluorescent, non-proteinaceous ligand (CuL) and we previously used fluorescence anisotropy to investigate the binding of this complex to PIC2 and SLC25A3 (11, 17, 21). Compared to PIC2, purified MIR1 showed limited interaction with the CuL complex (Fig. 1G). Thus, while the growth assays indicate that *MIR1* deletion can produce a Cu-dependent respiration defect at high Ag^+^ concentrations, our biochemical data suggest that MIR1 does not transport Cu. Therefore, both MIR1 and PIC2 transport phosphate but only PIC2 can transport Cu.

### Mitochondrial Cu and phosphate carriers duplicated early in the evolution of eukaryotes

It is not surprising that MCF proteins are present across all eukaryotes, given their fundamental roles in maintaining cellular physiology. In fact, we hypothesize that Cu transport to mitochondria was an important consideration in eukaryogenesis based on the central role of COX activity in the initial endosymbiosis (22). Conservation of this activity across diverse organisms may provide a phylogenetic signal with which to resolve residues involved in PIC2 and MIR1 substrate specificities. One evolutionary hypothesis is that because ancient proteomes were smaller, transporters in these organisms were generalists that gained specificity as a consequence of gene duplication and subsequent subfunctionalization (5, 6, 23, 24).

To provide evolutionary context for the existing experimental data, which has nearly all been collected from mammals and yeast, we performed phylogenetic analysis on PIC2/MIR1/MCF transporters from a broad range of eukaryotic lineages. We selected a set of 47 taxa for analysis that spanned the supergroups within the eukaryotic Tree of Life (eToL) (25) (Dataset S1). Only taxa with complete nuclear and mitochondrial genome sequences were included to accurately enumerate gene duplications and losses and ensure that apparent losses were not due to incomplete datasets. From these genomes, a total of 2,445 putative MCF family members were identified based on the presence of a mitochondrial carrier domain (PFAM domain PF00153). To distinguish PIC2-MIR1 orthologs from other members of the MCF family, phylogenetic trees were constructed using the MCF proteins from each taxon as well as the complete set of yeast and human MCF proteins. Candidate sequences that clustered with PIC2 or MIR1 were retained for further analyses (92 out of 2,445 MCF sequences).

Amino acid sequences of these potential Cu and/or phosphate transporting proteins were aligned and subsequently used to reconstruct the evolutionary history of PIC2-MIR1 orthologs across eukaryotes (Fig. 2). Of the 92 sequences, 47 clustered with *S. cerevisiae* PIC2 and are referred to as PIC2-like while 42 clustered with *S. cerevisiae* MIR1 and are defined as MIR1-like. The remaining three sequences were more closely related to PIC2-MIR1 than other MCFs but nonetheless fell outside of these two well supported clades.

**Figure 2.**
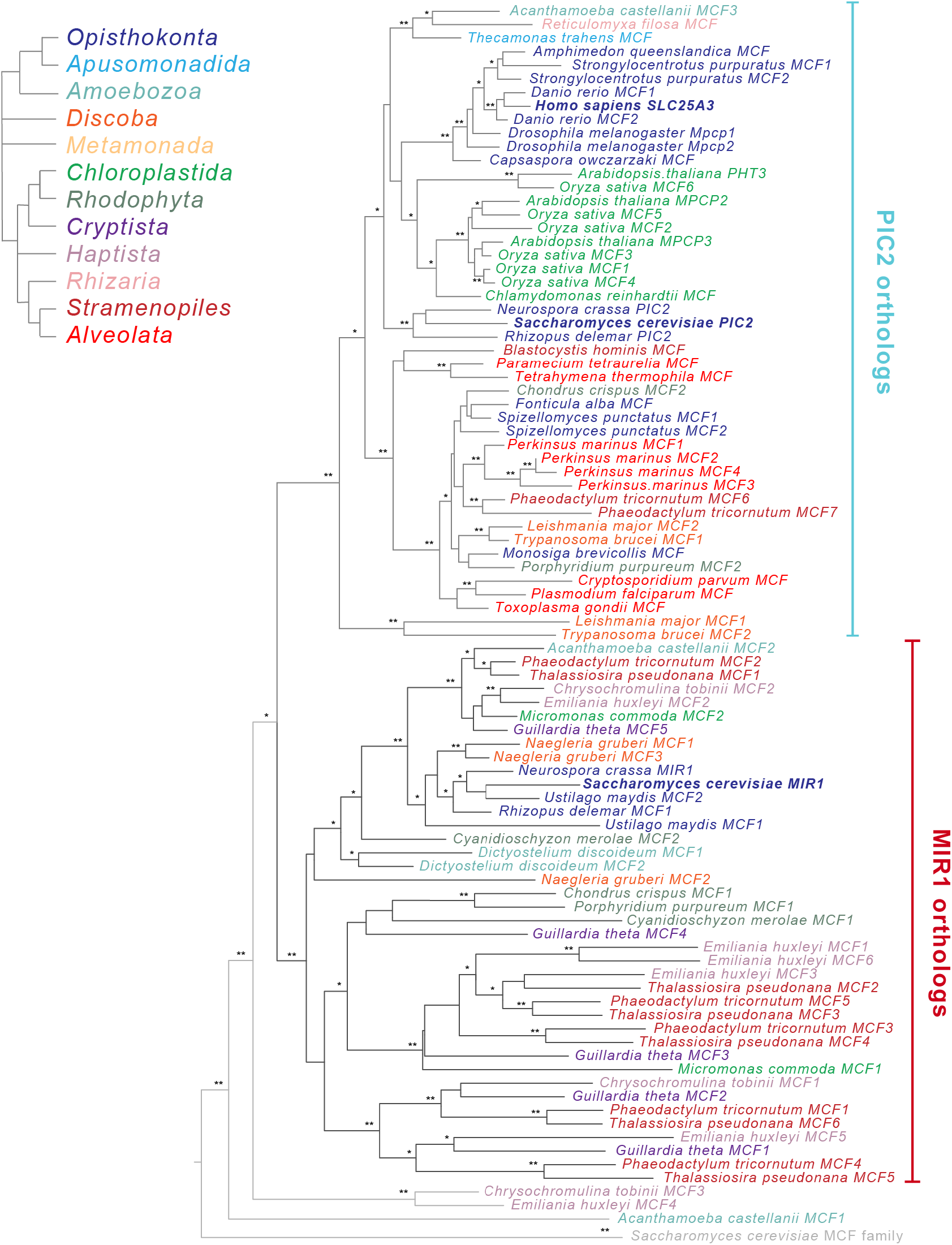
Phylogenetic analysis of the PIC2/MIR1 orthologs from 47 taxa reveals two major clades. Amino acid sequences of the eukaryotic MIR1/PIC2/SLC25A3 orthologs were aligned with the complete set of MCF proteins from *S. cerevisiae*. The maximum-likelihood tree shown was constructed in iQ-TREE using a general codon exchange matrix for nuclear genes with amino acid frequencies determined empirically from the data and seven rate categories (LG+F+R7). Support for the nodes was calculated using 1,000 replications and is indicated as follows: ** >95%; * >75%. Taxa names for the MIR1/PIC2/SLC25A3 sequences are color-coded according to the eToL supergroups as indicated; the *S. cerevisiae* MCF outgroup sequences (grey) have been collapsed to a single branch. Accession numbers for each of the sequences is available in Supplemental Dataset S1.

To estimate the timing of gene duplications and losses within the eukaryotes, we overlaid the presence and/or absence of PIC2-like or MIR1-like sequences onto the established eToL tree (Fig. 3A). Recent phylogenomic analyses indicate that extant eukaryotes form nine supergroups (25). Species from seven of these groups were included in this analysis: Amorphea, Discoba, Archaeplastida, TSAR (Telonemids, Stramenopiles, Alveolates, and Rhizaria), Haptista, Cryptista and Metamonada. Two additional groups, CRuMs (Collodictyonids, Rigifilida, and Mantamonas) and Hemimastigophora, were not included due to lack of complete nuclear genome sequences. PIC2-MIR1 orthologs were present in each taxon analyzed with the exception of those from Metamonads, which are anaerobic protists that secondarily lost mitochondria (26, 27), suggesting that the two paralogs were present within the last common eukaryotic ancestor (Fig. 3A).

**Figure 3.**
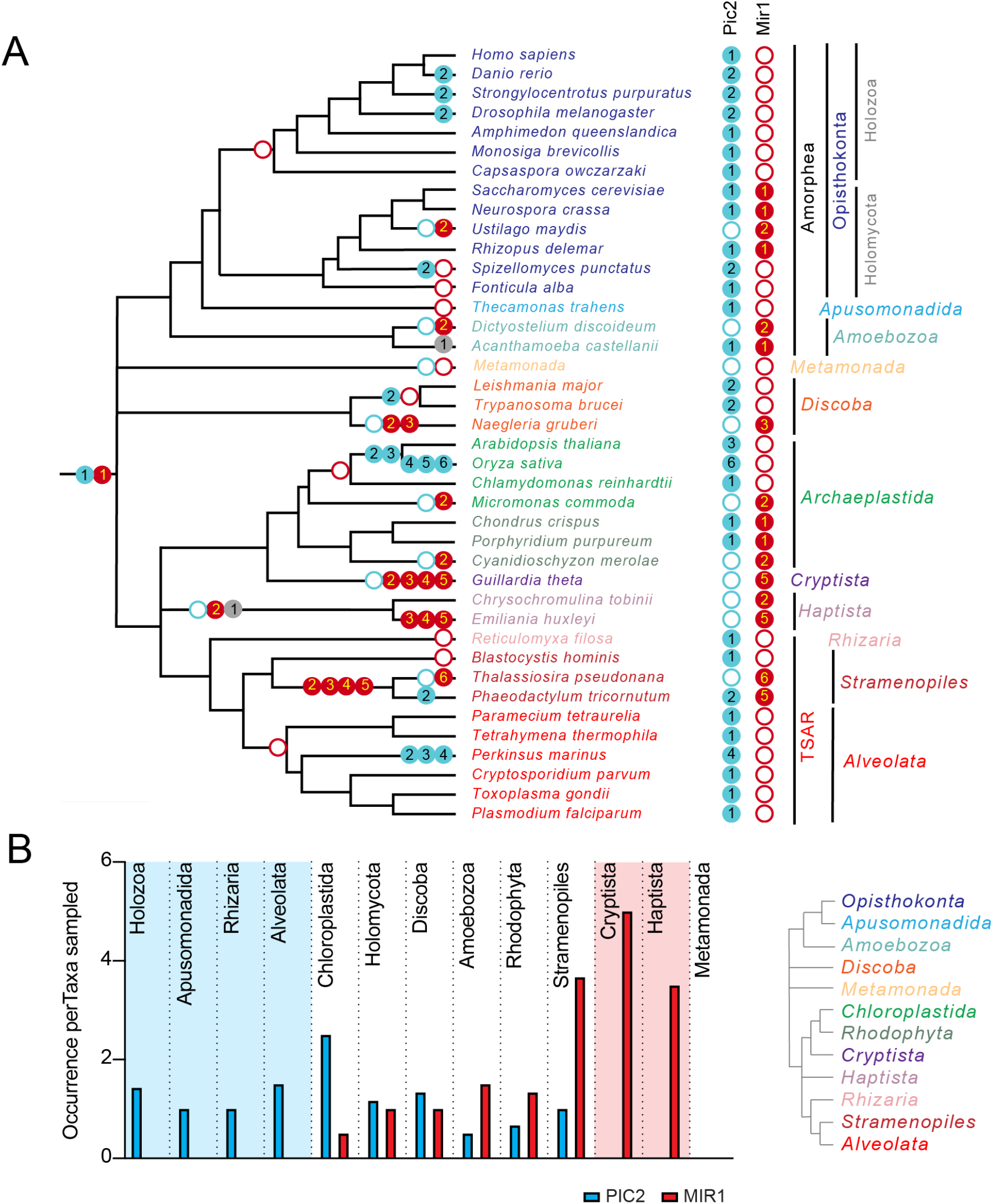
The PIC2/MIR1 family of MCF transporters is ancient within eukaryotes. *A)* Using the presence or absence of orthologs within the eukaryotic lineages, we inferred the evolutionary timings of gene duplications (solid circles) and losses (hollow circles) of the PIC2-like (blue), MIR1-like (red) and other (grey) sequences. *B)* The average number of PIC2 and MIR1 orthologs identified in the sampled taxa from eight of the nine eukaryotic supergroups.

Given the ancient origin of PIC2 and MIR1, we first analyzed the presence and absence of orthologs within Amorphea, which consists of the Opisthokonts (animals, fungi and yeast), Apusomonads and Amoebae (25). MIR1-like sequences are absent from Holozoan taxa with this lineage retaining only PIC2-like transporters (Fig. 3A, B). In contrast, the fungal lineages (Holomycota) exhibit more variability in the numbers of PIC2-like and MIR1-like sequences (Fig. 3B). Single orthologs of each type are present in *S. cerevisiae* and the closely related *Neurospora crassa*. The only Amorphea taxa that lost PIC2 are *Ustilago maydis* and *Dictyostelium discoideum* which both have a *MIR1* duplication. Outside the Amorphea, the gene copy number of the PIC2-MIR1 orthologs is more variable, which may reflect different evolutionary pressures on these transporters across lineages. Several lineages have lost either PIC2 or MIR1 and retained multiple copies of the remaining paralog (*e*.*g*., PIC2-like transporters within Chloroplastida and the alveolate *Perkinsus marinus* or the MIR1 duplications in Cryptista and Stramenopile lineages; Fig. 3). This raises the possibility that to compensate for the loss of the MIR1 transporter, PIC2 duplicated and convergently evolved additional substrate specificities. While there may be other constraints on this evolution, the loss of a PIC2 ortholog is always accompanied by duplication of the MIR1 ortholog. In contrast, a PIC2-like MCF is retained in all species that have a single PIC2-MIR1 ortholog, indicating that the loss of MIR1 does not always coincide with PIC2 duplication.

### Structural modeling of PIC2 suggests appropriate spatial organization of conserved residues that may coordinate Cu transport

We hypothesize that specific residues in PIC2*-*like proteins that confer the ability to transport Cu are absent in MIR-like proteins, while amino acids conserved across PIC2- and MIR-like proteins are required for both Cu and phosphate transport. To predict residues involved in substrate specificity we modeled the PIC2 sequence onto the c-state and m-state structures of the ADP/ATP carrier (8, 9) (Supplemental Fig. 1). Sequence conservation was calculated based on Shannon entropy using alignments of the PIC2-like sequences (Fig 4A, B)(Dataset S2). By integrating the structural models and phylogenetic analyses, we were able to visualize conserved residues as a surface representation (Fig. 4C, D, E). The PIC2-like orthologs show high conservation in the channel whereas alignment with the complete PIC2-MIR1 family reveals a smaller subset of conserved residues (Supplemental Fig. 2). This analysis also detects conserved patches extending into the IMS and outside the channel in the lipid bilayer that may be required for interactions with other components of the IM (Fig 4D, E).

**Figure 4:**
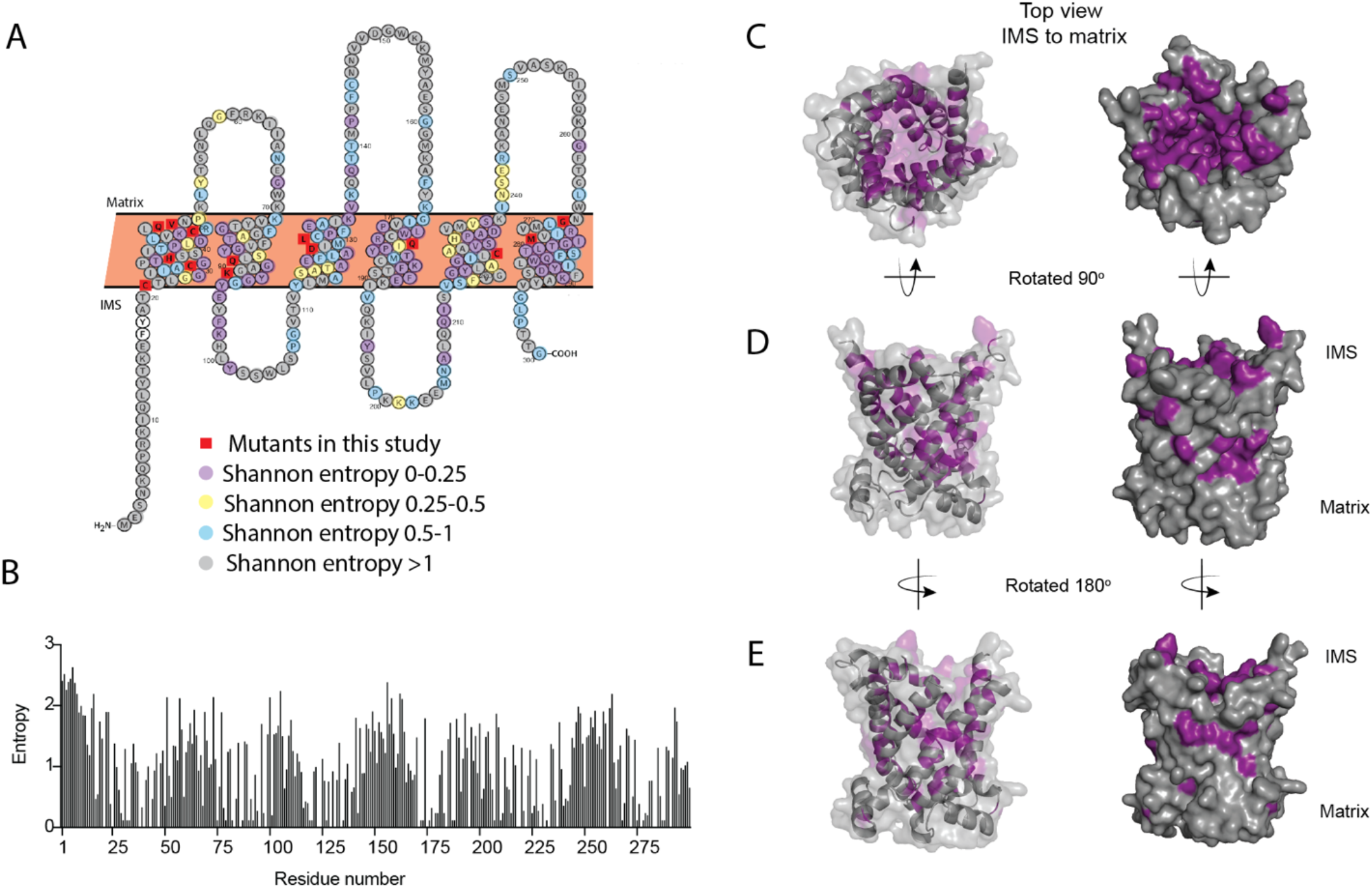
Conservation of residues in PIC2. *A)* A Protter representation of the PIC2 amino acid sequence was generated and colored based on Shannon entropy scores for conservation of a given residue. *B)* The Shannon entropy for each residue in PIC2 based on all sequences in the PIC2 specific clade (see Supplemental Dataset S2). *C)* Structure of PIC2 in the c-state viewed from the IMS side, with residues with >0.5 Shannon entropy highlighted in purple and all other residues colored grey. *D)* 90° rotation of the structure to view it from side and *E)* a 180° rotation to view it from the opposite side.

To identify residues for Cu transport, we initially focused on the well-established Cu-binding ligands Cys, His and Met. Analysis of the PIC2-MIR1 ortholog trees showed that histidine 33 (His33) (using the PIC2 numbering) is conserved in both the PIC2 and MIR1 clades (Fig. 5). Cysteine 29 is conserved in the PIC2 clade and most MIR1s but is replaced with Ala in the MIR1-like transporters from lineages with multiple duplications (*Emiliana huxleyi, Thalassiosira psuedonana*, and *Phaeodactylum tricornutum*). Cysteine 21 and Cys225 are strictly conserved among PIC2 orthologs, but not among MIR1 orthologs (Fig. 5). Cysteine 44 is conserved in the PIC2-like clade while MIR1-like orthologs have a conserved threonine in the equivalent position (Fig. 5). The PIC2-like transporters that lack Cys44 are the *P. marinus* duplications, one of two copies of PIC2 in *P. tricornutum* and the single copy of PIC2 in *N. crassa*. Analysis of the structural models revealed that Cys21, Cys29, Cys44 and His33 are positioned along one side of the channel (Supplemental Fig. 1) whereas Cys225 is on the opposite side of the channel. Cysteine 225 is positioned to interact with the peptide backbone of Cys182 (based on the alignments this residue is only a cysteine in *S. cerevisiae*), which faces away from the channel. Together, these data suggest that Cys21, Cys29, Cys44 and His33 may combine to form transient sites that bind Cu directly as it moves through the IM.

**Figure 5.**
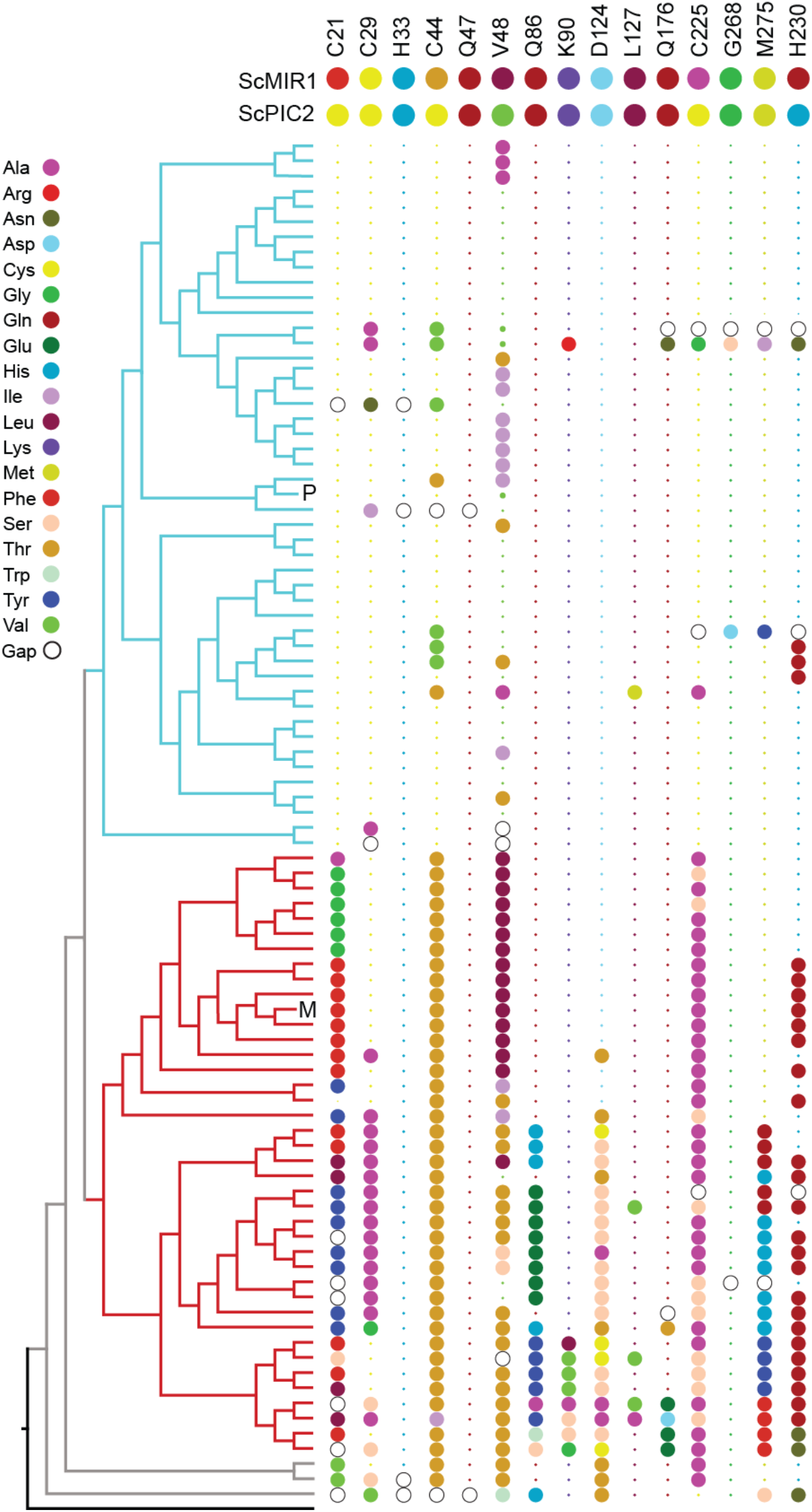
Conservation of selected residues in the PIC2/MIR1 family of transporters. The tree topology is identical to that shown in Figure 2. Amino acids are colored according to the key, and insertion/deletion events that lead to gaps within the alignment are indicated by the hollow circles. P indicates position of *S. cerevisiae* PIC2 and M indicates *S. cerevisiae* MIR1.

### Mutating structural elements and conserved contact points cause differential transport defects

To assess the functional importance of the Cys-His residues in Cu and/or phosphate transport we expressed PIC2 mutants in *L. lactis*. To assay Cu transport, we cultured each variant in media containing an Ag^+^ concentration that inhibited growth of *L. lactis* expressing wild-type PIC2 but not of cells harboring an empty vector (EV; Fig. 1A, Fig. 6). *L. lactis* expressing C21A, C29A, H33A, C44A and C225A PIC2 mutants displayed increased Ag^+^ resistance relative to *L. lactis* expressing wild-type PIC2 (all *P* < 0.012) (Fig. 6A), with the most resistance observed in the H33A mutant. However, these mutants also exhibited a growth defect relative to cells with an EV, suggesting that although transport is reduced residual activity is nonetheless present. Similarly, when Ag^+^ was replaced with AsO_4_^3-^to assess phosphate transport, *L. lactis* expressing each of the five PIC2 mutants displayed increased resistance to AsO_4_^3-^(Fig. 6B) suggesting that these mutations also limit its transport.

**Figure 6:**
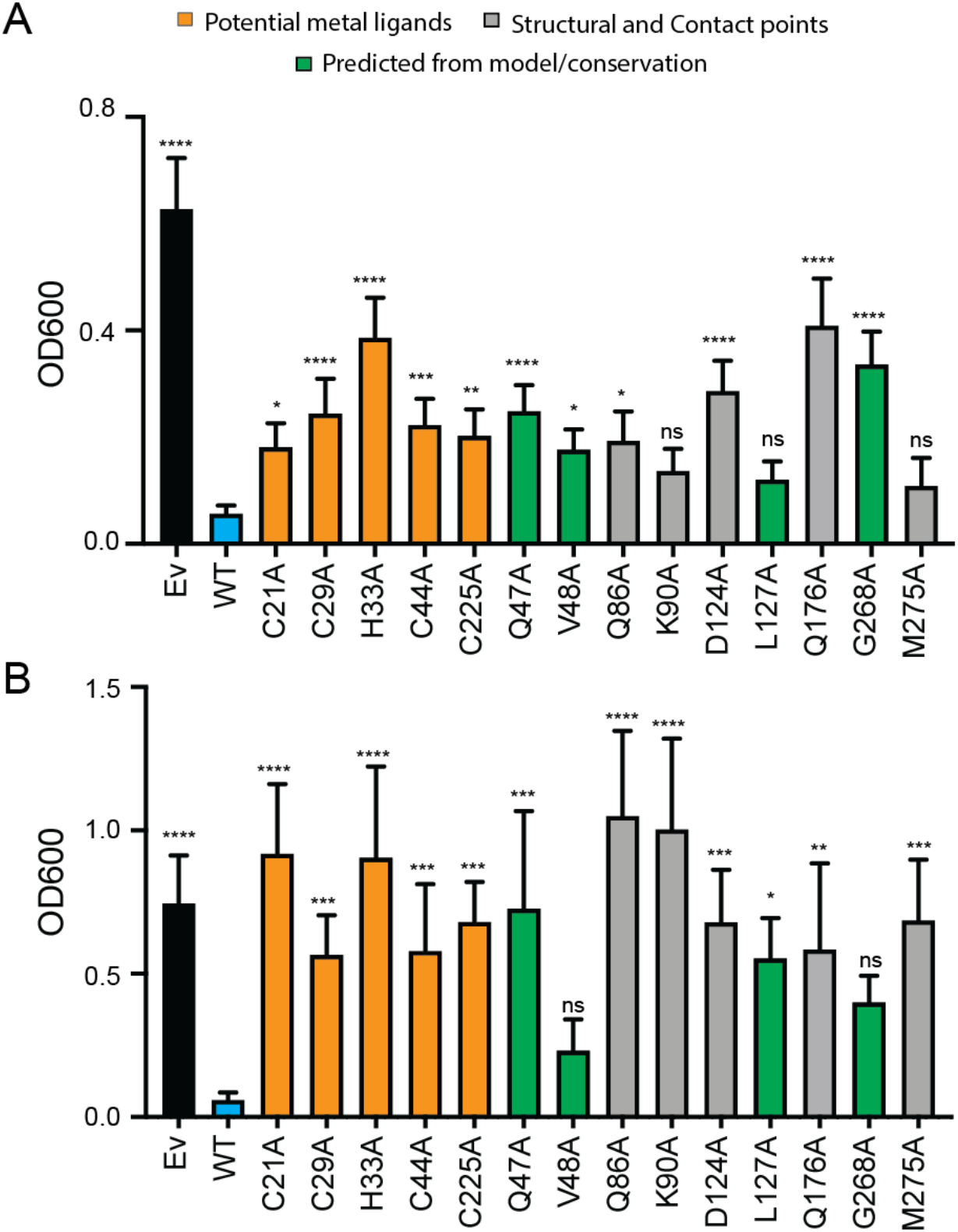
Expression of PIC2 and variants in *L. lactis*. *A)* Growth of *L. lactis* expressing EV, wild-type PIC2 (WT) or a given PIC2 variant in which each of the listed residues was converted to an alanine in Ag^+^ containing media. Each bar represents the median of 12 independent cultures with 95% confidence interval as error bars (*, P < 0.05, **, P<0.01, ***, P<0.001, ****, P<0.0001 based on one-way ANOVA relative to PIC2 wild-type control). The color of the bar indicates one of three major groupings; Cu-binding (orange), structural motifs or contact points (grey) and evolutionarily conserved and present in channel of the transporter (green). *B)* As described in *A)* except *L. lactis* strains were grown in AsO_4_^3-^containing media.

Computational analyses predict that *S. cerevisiae* MCF transporters have three contact sites for substrate binding (3). In PIC2, the proposed phosphate substrate contact points are Gln86 and Lys90 in transmembrane helix (TMH) 2, Gln176 in TMH4 and Met275 in TMH6 (Fig. 4) (3, 4, 8). These residues are largely conserved in both the PIC2-like and MIR1-like clades (Fig. 5), as is expected for transporters that share a substrate. We mutated each of these residues to alanine and assessed transport activity as described above. When expressed in *L. lactis*, the Gln86Ala and Gln176Ala mutants were more resistant to Ag^+^ than wild-type PIC2 (Fig. 6A) but less resistant than cells expressing EV. In contrast, the Lys90Ala and Met275Ala mutants exhibited comparable Ag^+^ sensitivity to wild-type PIC2 (p>0.05), suggesting that these substitutions do not affect Cu transport (Fig. 6A). The addition of AsO_4_^3−^to the media only inhibited the growth of cells expressing wild-type PIC2; cells expressing Gln86Ala, Lys90Ala, Gln176Ala and Met275Ala all grew at similar rates as cells expressing the EV (Fig. 6B).

Finally, we interrogated the functional significance of a subset of residues that were selected based on sequence conservation and our structural model; Gln47, Val48, Asp124, Leu127 and Gly268 (Fig. 4A, Supplemental Fig. 1). With very few exceptions, Gln47 is conserved among eukaryotic PIC2-MIR1 orthologs (Fig 4A, B and Fig. 5). Val48 it is part of a group of residues that appear to close the channel in the c-state (Supplemental Fig. 1). Asp124 interacts with Gln176 (Supplemental Fig. 1) and is conserved amongst all PIC2-like orthologs and those transporters most closely related to yeast MIR1 (Fig. 5). Leu127 is conserved in all orthologs and interacts with Gln86 (Fig 4A, B, Fig. 5, Supplemental Fig. 1). Gly268 is almost invariant throughout the evolution of this protein family (Fig 4A, Fig. 5). The Gln47Ala, Val48Ala and Asp124Ala PIC2 mutants expressed in *L. lactis* were more resistant to Ag^+^ than wild-type PIC2 (Fig. 6A) but less resistant than cells expressing EV, suggesting they also harbored residual Cu transport activity. When expressed in *L. lactis*, the Leu127Ala PIC2 variant showed equivalent susceptibility to Ag^+^ as the wild-type PIC2 but was resistant to AsO_4_^3-^(Fig. 6A, B) indicating that this single substitution interferes with phosphate transport but does not prevent Cu transport. Substituting alanine for valine at position 48 resulted in a significant difference in Ag^+^ resistance but mildly altered AsO_4_^3-^resistance (Fig. 6A, B). Finally, expression of the Gly268Ala variant resulted in resistance to Ag^+^ and AsO_4_^3-^, suggesting that this mutation disrupts the ability to transport both substrates (Fig. 6A, B). We also tested a series of mutants that exchanged the residues found in yeast PIC2 and mammalian SLC25A3 with those found in MIR1. Conversion of the PIC2 residues Ser102, Tyr156, Thr180, Gln138, Glu242 and Val191 to the equivalent residues in MIR1 did not affect the ability to transport Ag^+^ (Supplemental Fig. 3). Collectively, the data from the *L. lactis* assays show we can mutate individual residues that impair the transport of either Cu or phosphate or both.

### Mitochondrial Cu transport is compromised in a Leu175 mutant of SLC25A3

Based on the His33 and Leu127 PIC2 mutant data from *L. lactis*, we investigated the transport activity of the equivalent variants in murine SLC25A3 (His75 and Leu175). Consistent with the failure of the His33Ala PIC2 mutant to transport Ag^+^ or AsO_4_^3-^in *L. lactis*, expression of the equivalent His75Ala SLC25A3 variant in immortalized mouse embryonic fibroblasts (MEFs) with floxed (WT) or collapsed (KO) *Slc25a3* alleles did not rescue the COX deficiency of the KO cells (Fig. 7A, B). Conversely, expression of the Leu175Ala SLC25A3 variant was able to reverse the COX defect (Fig. 7B). Immunoblot analysis showed that the Leu175Ala mutant was present in mitochondria and increased steady-state COX1 levels (Fig. 7C). Consistent with our previous studies using a mitochondrially-targeted Cu sensor (17, 28), we found that total mitochondrial Cu content was significantly reduced in KO MEFs and increased in KO MEFs expressing the Leu175Ala variant (Fig. 7D).

**Figure 7:**
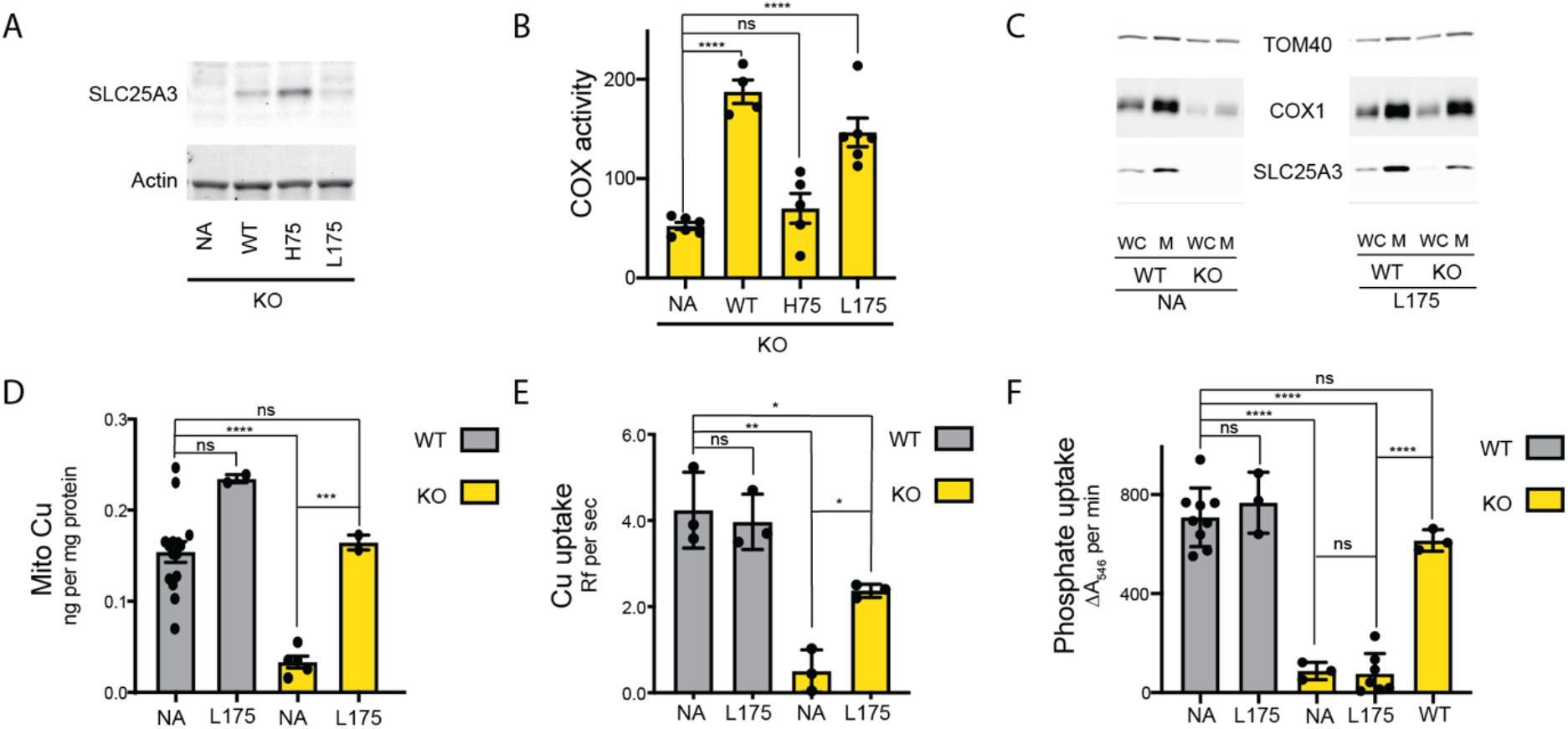
The SLC25A3 L175A variant restores mitochondrial Cu levels and rescues the COX deficiency in KO MEFs. *A)* Immunoblot analysis of SLC25A3 abundance in *Slc25a3 KO* MEFs alone or those transduced with wild-type SLC25A3 (WT), a His75Ala variant (H75) or a Leu175Ala variant (L175). Actin served as an internal loading control. *B)* COX activity in KO MEFs alone (n= 6) or transduced with WT SLC25A3 (n = 4), a His75Ala variant (H75) (n=5) or a Leu175Ala variant (L175) (n=6). ns, P>0.05, ****, and p < 0.0001 based on a one-way ANOVA. *C)* Immunoblot analysis of SLC25A3, TOM40 and COX1 abundance in whole cells (WC) or isolated mitochondrial (M) from WT or KO MEFs alone (NA) or transduced with the SLC25A3 Leu175Ala variant (L175). *D)* Total Cu levels in mitochondria from WT or KO cells as in *C)*, determined by ICP-OES. *E)* Cu uptake in mitochondrially derived liposomes created by the membranes of mitochondria in *C)* with additional lipids. Liposomes contain Phen green to monitor the uptake of Cu. *F)* Mitochondrial swelling rate in presence of phosphate as a measure of phosphate uptake.

Reconstitution of MCF proteins in liposomes has been used extensively to assess substrate transport and specificity (14, 29-33). Liposomes created from mitochondrial membranes of WT but not KO MEFs were able to transport Cu (Fig 7E). The Cu transport defect in KO-derived liposomes was reversed upon expression of the Leu175Ala variant (Fig 7E). To assess phosphate uptake, mitochondrial swelling in the presence of phosphate was measured (12, 18). Intact mitochondria isolated from KO cells had a phosphate uptake defect compared to WT that was rescued by expressing WT SLC25A3 but not the Leu175Ala variant (Fig. 7F). Taken together, these data show that the Leu175Ala mutant is able to transport Cu but not phosphate in mitochondria and that this Cu transport activity is sufficient to rescue COX activity.

## Discussion

The mechanisms that mediate MCF transporter specificity remain largely unknown. While individual studies have investigated deficiencies in the transport of one substrate, few have assessed substrate promiscuity. Here, we directly addressed this issue by focusing on Cu and phosphate transport which, in mammals, is mediated by the single MCF transporter SLC25A3. Multiple studies clearly connect SLC25A3 to phosphate transport and mutations in *SLC25A3* lead to skeletal muscle myopathy and heart disease in humans (17, 18, 34-37) and cardiac hypertrophy in mice (18). *Slc25a3* knockout MEFs derived from the heart-specific *Slc25a3* knockout mouse exhibit clear COX and SOD1 defects that can be rescued by overexpression of a *Slc25a3* cDNA or addition of Cu (17). These data are complemented by *in vitro* Cu transport by purified SLC25A3 in liposomes and by Ag^+^ growth phenotypes associated with its expression in *L. lactis* (17). The data presented in this study provide the first experimental evidence of a missense mutation that separates Cu and phosphate transport, and firmly establish that physiological defects in COX and SOD1 are due to Cu transport and not secondary effects resulting from decreased phosphate transport.

### Evolutionary history of mitochondrial Cu-phosphate transporters

Our evolutionary analyses of the Cu-phosphate transporters were prompted by the observation that *S. cerevisiae* PIC2 and MIR1 exhibit substrate specificity, whereas the mammalian ortholog SLC25A3 is responsible for the transport of both Cu and phosphate. Selection on genes with multiple functions can constrain diversity to avoid negative effects associated with losing one of these functions. Therefore, gene duplications serve as important sources for evolutionary selection and refinement. Resulting duplications can be retained for the original function, specialized for new functions, refined to enhance an existing function or allow for increased expression by gene dosage; if none of these occur, the duplicate gene is lost (38-44). In *S. cerevisiae*, PIC2 and MIR1 are partially redundant for phosphate transport (12). However, mutation of *MIR1* in *S. cerevisiae* is sufficient to produce phosphate-related phenotypes suggesting that, under most conditions, the ability of PIC2 to transport phosphate is unable to compensate for loss of MIR1 function (12, 17). Instead, the *PIC2* sequence appears to be optimized for Cu transport. Similarly, we show here that MIR1 lacks clear Cu transport activity even though *mir1Δ* yeast exhibit increased susceptibility to Cu restriction compared to WT cells. Our phylogenetic analyses of *PIC2* and *MIR1* sequences suggest that the gene duplication that created these two orthologs was an ancient event, and that evolutionary interplay between these two substrate specificities may have occurred multiple times throughout eukaryotic evolution.

The loss of *MIR1* has occurred multiple times in eukaryotes, an event that is likely facilitated by the dual specificity of PIC2. *SLC25A3* is essential in mammals as the homozygous deletion is embryonic lethal. While mammals do express two SLC25A3 isoforms, isoform A is expressed primarily in heart and skeletal muscle whereas isoform B is expressed in all tissues (14, 18, 34). Therefore, it is unlikely that the isoforms provide the functional redundancy that would be afforded via gene duplication or retention of *MIR1*.

### Understanding Cu transport

The Leu175Ala mutation in SLC25A3 that separated Cu and phosphate transport fully restores COX activity and mitochondrial Cu levels without rescuing phosphate transport. This finding confirms that the COX defect in mutant cells is due to defective Cu transport, rather than reduced phosphate levels. Further, our data suggest that compromising the phosphate transport function of PIC2 is easier than inactivating its Cu transport function. Mutations in a series of cysteine and histidine residues lining the channel of the c-state model decrease, but do not eliminate, Cu transport. The PIC2 structural model indicates that the Cys29 and His33 would be the most likely location to form a Cu-binding site. The cysteine positioned above that site (residue 21) may help recruit Cu from the IMS and present it to Cys29-His33. In the m-state model, the Cys29-His33 proximity is maintained and the next potential ligand, Cys44, is exposed allowing for potential relocation of the Cu. PIC2 is able to transport Cu supplied in multiple forms, including a ligated form known as CuL that is present in the mitochondrial matrix (11). The CuL complex is negatively charged, suggesting that positively charged or hydrogen-bond donor residues within the channel may stabilize this interaction, including those that participate in phosphate transport (Gln86 and Lys90) (45). Additionally, 1D and 2D heteronuclear NMR analysis of the purified CuL shows the presence of a substituted benzene ring structure consistent with its fluorescent properties (Supplemental Fig. 4). In the m-state, the aromatic ring of the side chain of Tyr83 comes between the cysteine and histidine. This structure could mimic a CuL-bound state (from the c-state) and the movement of the side chain could facilitate the release of the complex from the Cys29-His33 site towards the matrix (Supplemental Fig. 4). The spatial arrangement of these residues may allow for either CuL binding and subsequent release of Cu or facilitate transport of the intact CuL complex. The intact transport may be expected as this is the major form of Cu in mitochondria under normal conditions. In addition, the anionic nature of the CuL complex may explain some of the promiscuity between Cu and phosphate as substrates of the same carrier.

Our phylogenetic analysis revealed nine taxa that lack a PIC2-like ortholog yet have COX. Each of these taxa have multiple *MIR1*-like transporters (*Guillardia theta, Thalassiosira pseudonana, Emiliania huxleyi, Dictyostelium discoideum, Ustilago maydis, Cyanidioschyzon merolae, Chrysochromulina tobinii, Micromonas commoda*, and *Naegleria gruberi*). Alignment of these paralogs identified residues that are present in at least one of the duplicates and are shared with PIC2 (Fig. 5 and Supplemental Fig. 5). This analysis may highlight the variants that have allowed MIR1 to secondarily gain Cu transport activity. One consistent difference is a histidine found in PIC2 orthologs versus a glutamine found in MIR1 orthologs at position 230 (numbering for PIC2). Both of these side chains stabilize the conformation of a possible cardiolipin binding site, by hydrogen bonding to peptide carbonyl oxygens. Additional experiments will be required to determine if this substitution affects substrate selectivity.

In the taxa with multiple MIR1-like paralogs that lack PIC2-like transporters (*i*.*e*., *E. huxleyi, G. theta* and *U. maydis*), we also observe multiple changes in the residues studied here (Supplemental Fig 5). We favor a hypothesis in which *MIR1* duplication is a response to overcome the loss of PIC2. However, this requires further investigation and an acknowledgement that other MCF transporters may have also acquired Cu transport activity. Indeed, in yeast we have shown that the MCF family member MRS3 serves as a secondary importer of mitochondrial Cu (21). MRS3 is known as an iron transporter, but transport of Cu by MRS3 and its orthologs has been reported in studies using mitochondrially derived vesicles from yeast and plants and in a reconstituted assay system (46-49). We did not compare the presence or absence of MRS3 orthologs in these taxa.

### Understanding phosphate transport

Our biochemical data suggest that Lys90 and Leu127 are important for phosphate transport but dispensable for Cu transport in *L. lactis*. The proposed mechanism of transport for MCF based on the comparison of the c- and m-states of the ADP-ATP carrier suggests that even-numbered helices shift to allow transport/transition to the opposite state (4, 8). The PIC2 structural model shows that Leu127 is on helix 3 adjacent to a proline that kinks helix 3, thereby altering helix-helix packing interactions with helix 2 (Fig. 8). The Leu127 side chain interacts with the peptide backbone between Leu85 (Met in SLC25A3) and Gln86, a “knobs into holes” interaction. We hypothesize that helix 2 reorients in the alanine substitution mutant to have stronger van der Waals contact, especially in the vicinity of Gln86. In the c-state, this change could shift the side chains of Gln86 and Lys90 to a conformation that disrupts a phosphate binding site (Fig. 8).

**Figure 8:**
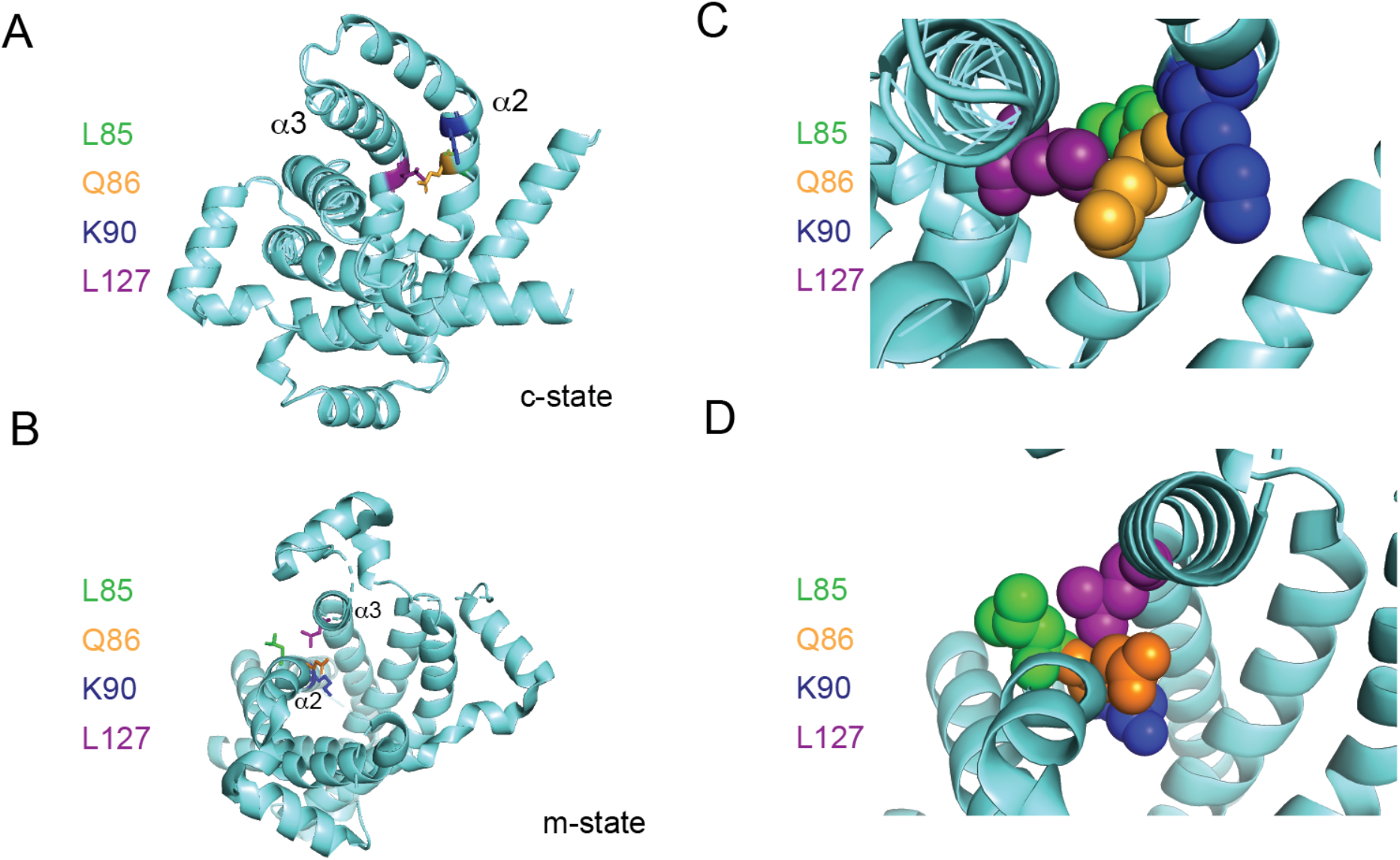
Positioning of Leu127 relative to adjacent residues on helix 2. Ribbon diagrams of PIC2 *A)* c-state and *B)* m-state structures. The polypeptide is shown as a ribbon trace (aquamarine), the side chains as stick models. The Leu127 is colored purple to distinguish it from the adjacent Leu85 (green), Gln86 (orange) and Lys90 (blue) residues on helix 2 (*α*2). Enlargement of the Leu127 interaction with the surrounding residues shown as spheres in *C)* c-state and *D)* m-state.

Cu transport requires the formation of transient covalent bonds between the metal and ligands during transport, whereas phosphate transport relies on hydrogen bonding and salt bridges. These requirements may account for the fact that multiple mutations were able to inhibit the ability of PIC2 to transport phosphate. Other site-directed mutational studies of MIR1 have identified multiple residues that are required for phosphate transport (50-54), including His33, Thr44 and Lys90 (using PIC2 numbering). Consistent with these earlier studies, we observe decreased phosphate transport when mutating the corresponding residues in PIC2. In fact, previous studies of MIR1 function showed that mutation of Thr44 to cysteine partially inactivated phosphate transport (54). This cysteine/threonine is clearly demarcated at the node between PIC2 and MIR1 clades, suggesting that it may be a critical change that weakened, but did not eliminate, phosphate transport in PIC2-like transporters (Fig. 5). Three lineages (*O. sativa, S. punctatus* and *P. marinus*) lack MIR1-like transporters and have multiple PIC2-like transporters. In the case of rice, this could simply be due to the polyploid nature of its genome. In the chytrid *S. punctatus*, it could suggest that duplication enhances gene dosage. That is, additional copies compensate for less efficient phosphate transport. In contrast, the duplicated genes in *P. marinus* have undergone several notable changes; one variant has a large carboxy terminal truncation, 3 of the 4 variants have valine replacing cysteine at position 44 (as noted above from previous studies threonine at this position is optimal for phosphate transport) and histidine at position 230 is replaced by the glutamine that is found in more phosphate-selective transporters. These changes and gene dosage may be sufficient to overcome the loss of a MIR1-like transporter. Testing these hypotheses will require *in vitro* expression of multiple transporters to assess substrate selection.

## Conclusions

Mitochondria function as a metabolic hub that controls physiology and disease by balancing the concentrations of multiple metabolites and essential elements (10, 55). The MCF proteins are a critical piece in regulating the import and export of these substrates (1, 2) and have been duplicated and specialized over evolutionary time to selectivity recognize and transport highly similar substrates. However, gene duplication has allowed for the retention of some carriers with multiple substrates. The evolutionary relationships among these carriers reveal aspects of transport mechanisms and the physiological demands of the organism. Our analysis of the Cu-phosphate MCF transporters shows that organisms deploy multiple strategies to recruit these substrates. We cannot determine a single characteristic that indicates an advantage or disadvantage of either strategy, as unique patterns appear nested in different lineages. Metal transport to the mitochondrial matrix is required for Fe-S cluster assembly and COX assembly. Perhaps metal substrates are sufficiently simple that multiple MCFs are capable of transport. However, given the fatal disorders that result from too much or too little Cu or iron it is unlikely that their transport is left to chance (56). Cu storage in the mitochondrial matrix may have evolved as a mechanism to ensure Cu availability for COX assembly in an early endosymbiont that was subsequently retained during eukaryogenesis (22). Additional roles for Cu in the matrix remain to be determined. The recent discoveries that mitochondrial Cu can induce cell death through a pathway coined cuproptosis (57), disrupt essential processes such as Fe-S assembly (58, 59) and alter the stability of SOD1 in the cytosol (17) collectively suggest that understanding the physiological consequences of disrupting this Cu pool and its homoeostasis remains an important area of future research.

## Methods

### Phylogenetic analysis

To delineate the evolutionary histories of the PIC2/MIR1 orthologs, 47 species were chosen that span the eukaryotic supergroups defined here. For each of these species, complete nuclear genome assemblies and protein predictions are available from NCBI (Supplemental Table 1). MCF orthologs were identified using HMMER (60) to detect sequences containing the Mitochondrial Carrier (MC) domain (PFAM PF00153). Redundant sequences and transcript variants were eliminated using CD-Hit with a threshold of 0.9 (61).

To distinguish PIC2/MIR1 orthologs from other members of the MCF family, phylogenetic trees were built using the MC domain-containing proteins from each organism as well as the complete set of MCF proteins from *Homo sapiens* and *Saccharomyces cerevisiae*. Amino acid sequences were aligned in MEGA X (62) using ClustalW with default parameters. Neighbor-joining trees were generated using a Poisson substitution model, uniform substitution rates among sites, and pairwise gap deletion. Support values were determined using 1,000 bootstrap replicates.

Amino acid sequences of the eukaryotic MIR1/PIC2 orthologs were aligned with 32 *S. cerevisiae* MCF proteins using MUSCLE implemented in MEGA X. Phylogenetic analysis was performed using IQ-TREE version 2.0.3 (63). The optimal substitution model was selected using the IQ-TREE ModelFinder (64). A maximum likelihood tree was constructed using the LG+F+R7 model (a general codon exchange matrix for nuclear genes with amino acid frequencies determined empirically from the data and 7 rate categories). Support was calculated based on 1,000 replications using ultrafast bootstrap approximation (UFBoot2;(65)).

### Structural modelling

Sequence alignments between Pic2 and the ADP/ATP exchanger were used correctly place indels and ensure proper alignment of the key helices. Initial molecular models were generated using Swissmodel and subjected to careful analysis in Coot for side chain rotamer optimization, interatomic clashes and hydrogen bonding (66). Finally, the model atomic coordinates were energy minimized within the PHENIX suite (67). We modeled Pic2 based on the m-state and the c-state of the ADP/ATP exchangers deposited in the protein data bank (PDB:4C9G and PDB: 6GCI).

### Expression in Lactococcus lactis

*L. lactis* cells transformed with vector (pNZ8148 (MoBiTec)) alone or pNZ8148 carrying yeast MIR1, PIC2 or site directed PIC2 mutants were grown overnight at 30°C in M17 medium with 0.5% glucose and 10 µg/mL chloramphenicol. To determine Ag^+^ toxicity in *L. lactis* strains containing vector, MIR1, PIC2 or PIC2 mutants, cells were grown in a 96-well plate containing M17 medium plus 1 ng/mL nisin and increasing concentrations of Ag^+^ (0-250 µM) or AsO_4_^3-^(0-2.5mM). Controls containing M17 without nisin or M17 plus Ag^+^ or AsO_4_^3-^without nisin were included. Optical density at 600 nm was used to assess growth after 24 hours. Percent growth was quantified by comparing to the optical density of the same genotype in nisin alone.

### Elemental analysis

Samples were digested in 40% nitric acid by boiling for 1 hour in capped, acid washed tubes, diluted in ultra-pure, metal free water and analyzed by ICP-OES (Perkin Elmer, Optima 7300DV) versus acid washed blanks. Concentrations were determined from a standard curve constructed with serial dilutions of two commercially available mixed metal standards (Optima). Blanks of nitric acid with and without “metal-spikes” were analyzed to ensure reproducibility.

### Cell culture conditions

Clonal Slc25a3^FLOX/FLOX^ and Slc25a3^-/-^MEF lines were then isolated and maintained in high glucose DMEM containing sodium pyruvate, 50 µg/ml uridine, 0.1mM mercaptoethanol and 15% fetal bovine serum at 37oC at an atmosphere of 5% CO2 (17). Mouse Slc25a3-b cDNA was amplified from RNA and cloned into a Gateway-modified retroviral expression vector. The fidelity of this construct was confirmed by sequencing and retrovirus was produced with the Phoenix Amphotrophic packaging cell line and used to transduce MEFs.

### Immunoblot and activity assays

This study used monoclonal antibodies raised against TOM40 (ProteinTech 18409-1-AP), and COX1 (Abcam ab14734), and a rabbit polyclonal antibody raised against the KLH conjugated SLC25A3 peptide CRMQVDPQKYKGIFNGSVTLKED (Pacific Immunology). COX activity was determined by monitoring the decrease in absorbance at 550 nm of chemically reduced cytochrome *c* in the presence of whole cell or mitochondrial extracts (65). All activities were normalized to protein concentration then converted to percentage of maximum control value.

## Acknowledgements

We thank Ann Ashton, Shannon Baseke, Shelley Brannon, LaQuasha Jones, Kacie Oglesby, Christina Waples, Mary Wetzel (Auburn University undergraduate research students) for contributing to construction and initial testing of the different mutants in this study. This work is supported by a grant from National Institutes of Health [R01GM120211 to PAC and SCL]. PAC is supported by the Alabama Agricultural Experiment Station. KMB is supported by a grant from National Science Foundation [EF 2021886].

## Legends for Supplemental Figures and Tables

**Dataset S1** (Supplemental Table 1.xlxs). The accession numbers of sequences analyzed in this study.

**Dataset S2** (Supplemental Table 2.xlxs). The comparison of PIC2 and MIR1 showing entropy scores and conservation of residues.

**Fig. S1.**
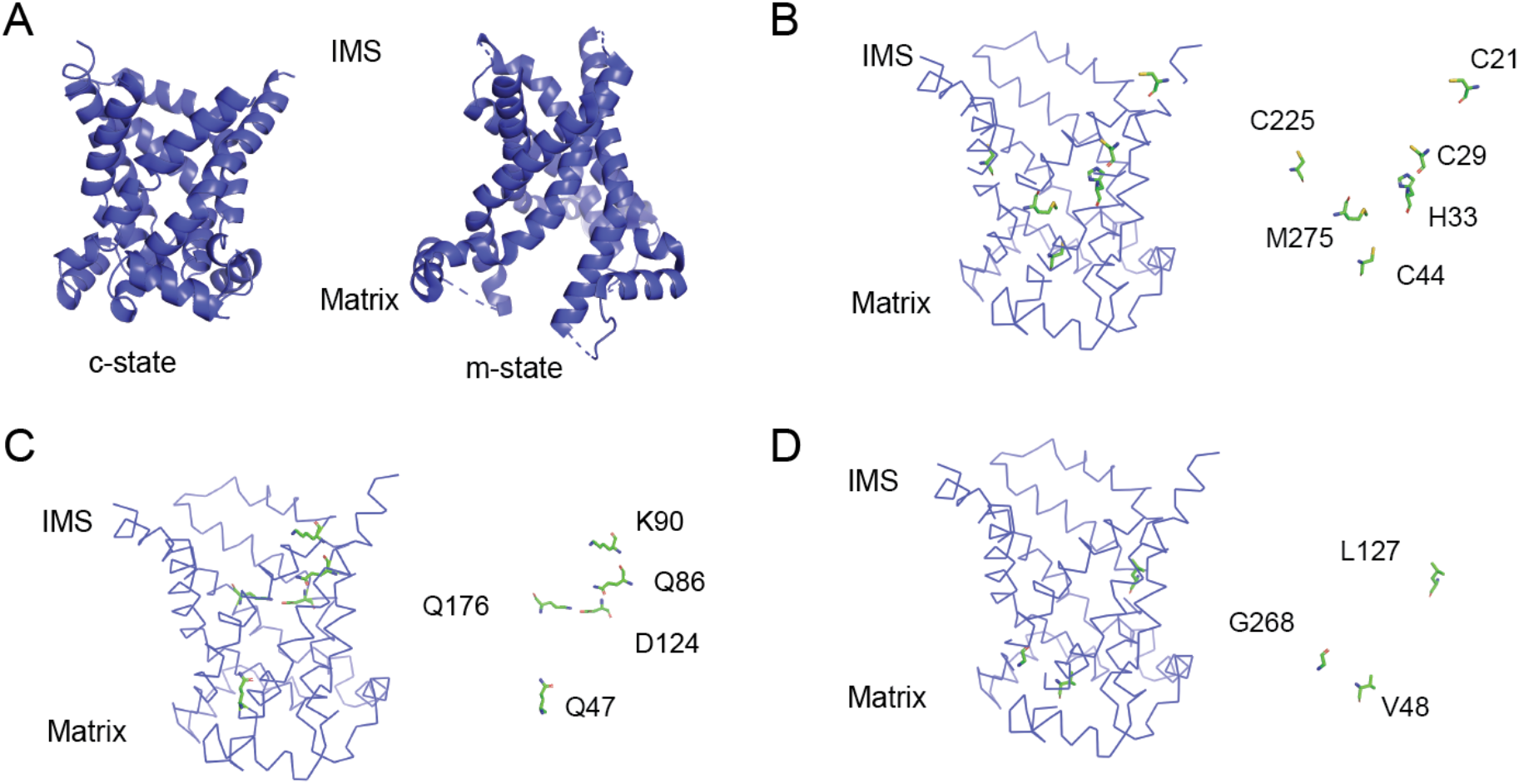
Structural models of PIC2 and representation of the residues mutated in Fig. 6. A) Cartoon ribbon structure of PIC2 modelled onto ADP-ATP carrier in the c-state and m-state. B) Model of the c-state of PIC2 with C21, C29, C44, C225, H33, M275 highlighted in sticks format and with the backbone cartoon representation removed. C) Model of the c-state of PIC2 with K90, Q47, Q86, D124, Q176 highlighted in sticks format and with the backbone cartoon representation removed. D) Model of the c-state of PIC2 with V48, L127, G268 residues highlighted in sticks format and with the backbone cartoon representation removed.

**Fig. S2.**
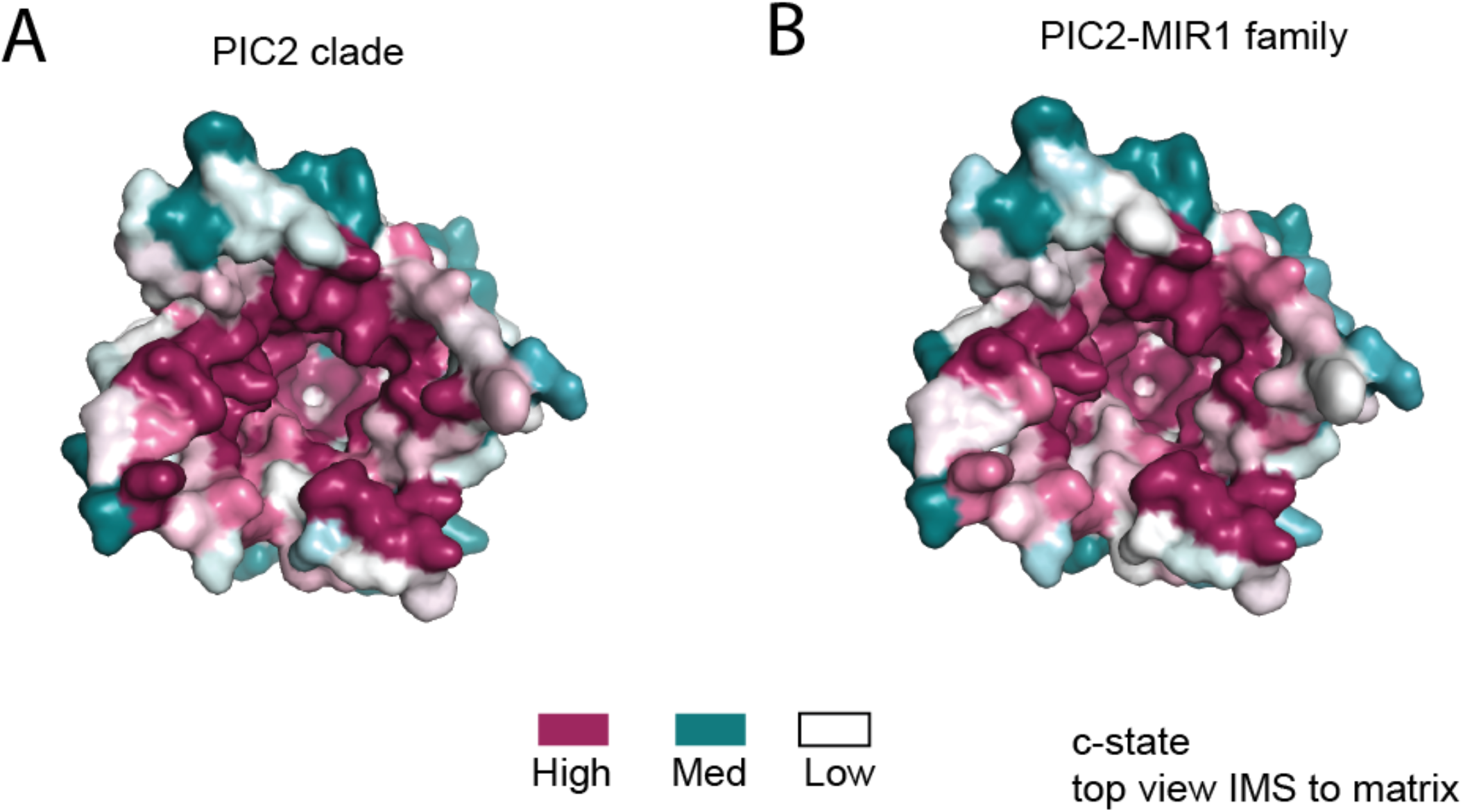
Conservation surface of PIC2 viewed from the IMS. The c-state model of PIC2 with surface representation colored based on in the conservation PIC2 clade as defined in Fig. 3 or based on the complete PIC2-MIR1 family.

**Fig. S3.**
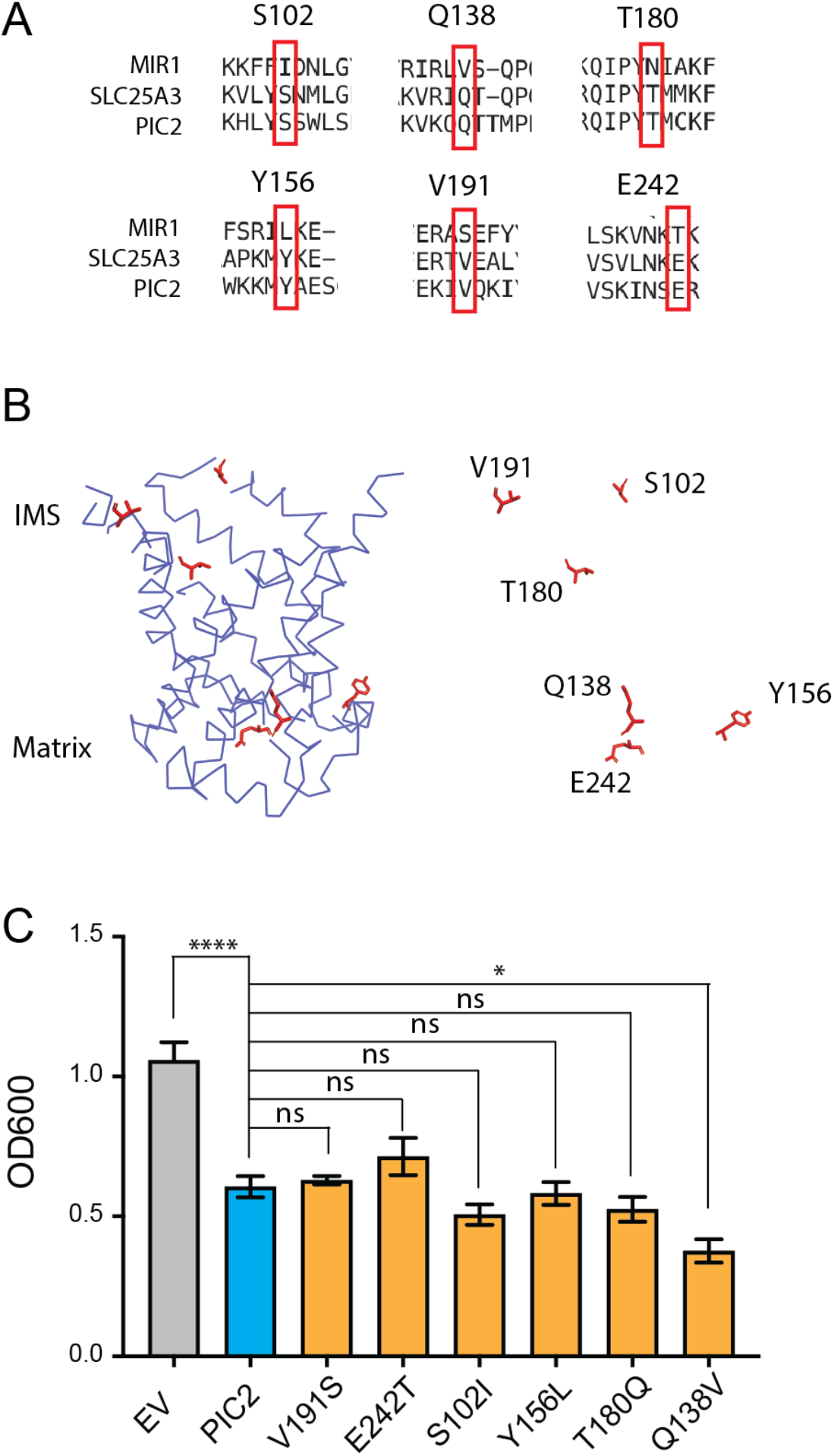
Substitution of PIC2 residues for MIR1 residues. A) Alignment of amino acid sequences from MIR1, PIC2 and SLC25A3 B) Model of the c-state of PIC2 with residues mutated highlighted in sticks format and with the backbone cartoon representation removed C) Growth of L. lactis expressing EV, wild-type PIC2 (WT) or a given PIC2 variant in which each of the listed residues was converted to an alanine in Ag+ containing media. Each bar represents the median of 6 independent cultures with 95% confidence interval as error bars (*, P < 0.05, **, P<0.01, ***, P<0.001, ****, P<0.0001 based on one-way ANOVA relative to PIC2 wild-type control).

**Fig. S4.**
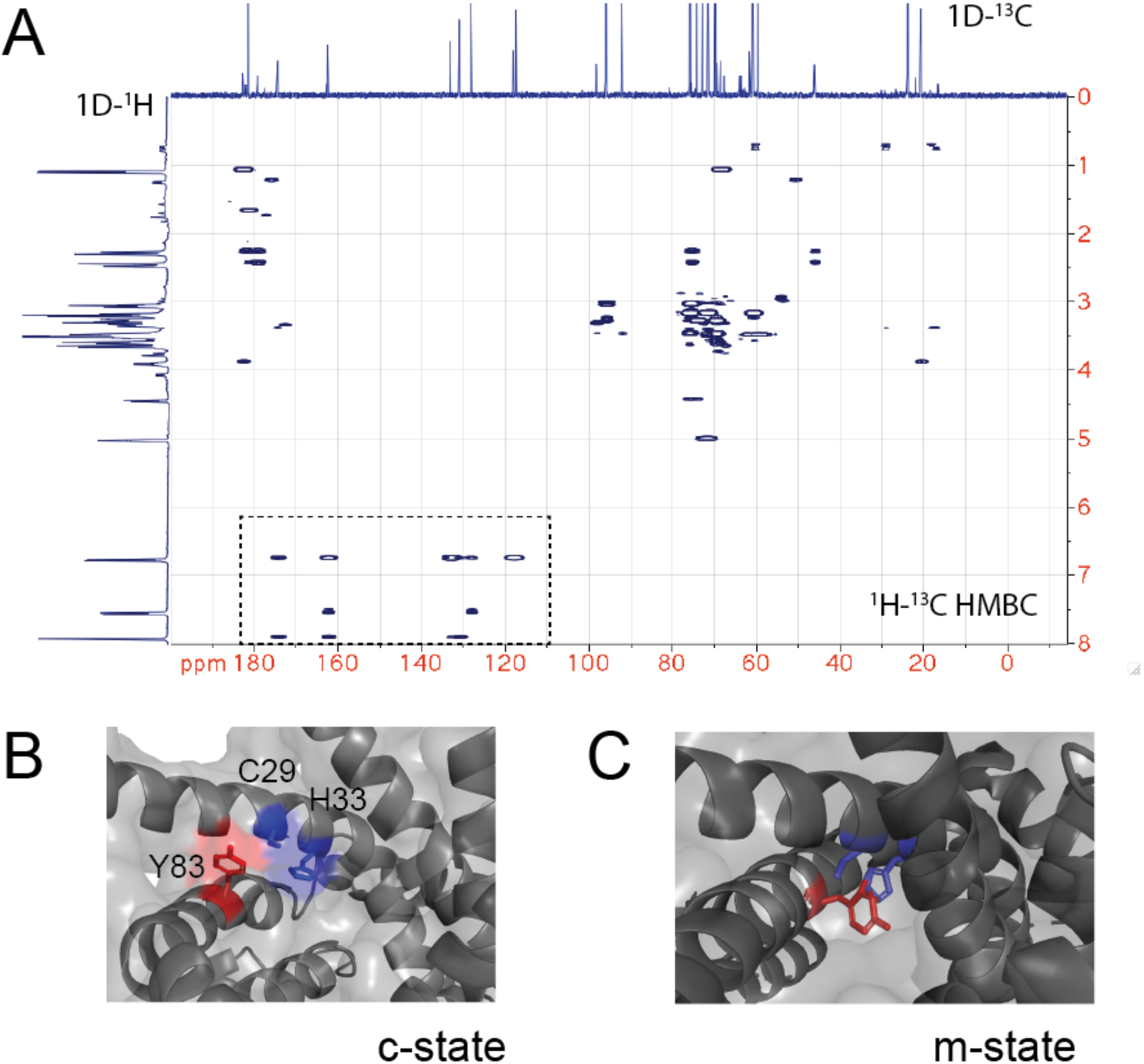
NMR of the CuL and role of Y83 in interactions with the proposed C29-H33 binding site A) ^1^H-^13^C HMBC spectrum of the purified CuL complex. The 1D ^1^H and ^13^C spectrum are shown. The box highlights the signals consistent with a benzene ring in the CuL. B) Enlargement of the C29-H33 region of c-state with C29, H33 and Y83 shown in sticks. C) Enlargement of the C29-H33 region of m-state with C29, H33 and Y83 shown in sticks with Y83 between the C29-H33 “replacing/occluding” a potential site for CuL binding.

**Fig. S5.**
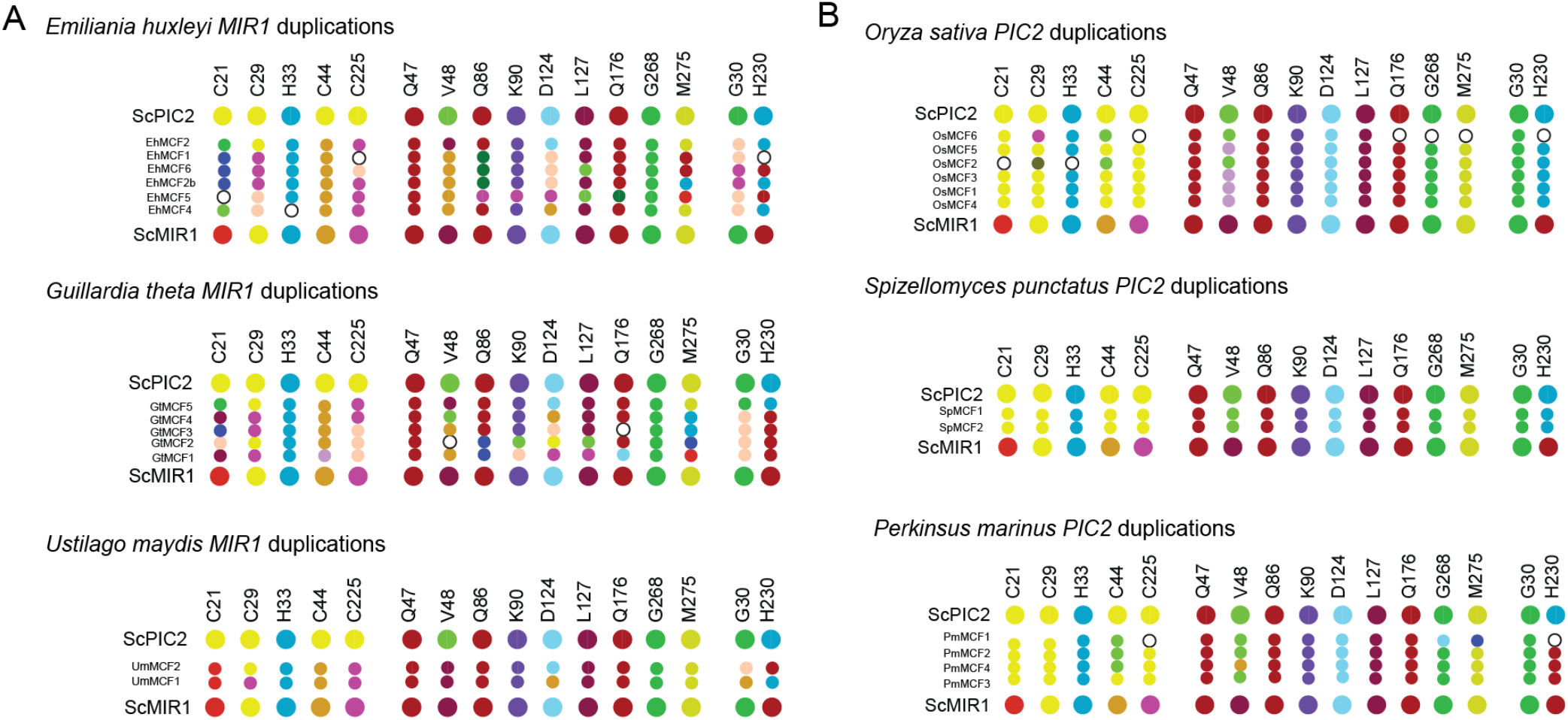
Map of the residues found in duplications. A) Graphical representation of the residues in organisms which have duplicated MIR1 and lack PIC2 B) Graphical representation of the residues in organisms that have duplicated PIC2 and lack MIR1.

